# Exploring the context of diacidic motif D-E as a signal for unconventional protein secretion in eukaryotic proteins

**DOI:** 10.1101/250076

**Authors:** Sreedevi Padmanabhan, Malay Ranjan Biswal, Ravi Manjithaya, Meher K. Prakash

## Abstract

Unconventional protein secretion (UPS) is an important phenomenon with fundamental implications to cargo export. How eukaryotic proteins transported by UPS are recognized without a conventional signal peptide has been an open question. It was recently observed that a diacidic amino acid motif (ASP-GLU or DE) is necessary for the secretion of superoxide dismutase 1 (SOD1) from yeast under nutrient starvation. Taking cue from this discovery, we explore the hypothesis of whether the diacidic motif DE, which can occur fairly ubiquitously, along with its context, can be a generic signal for unconventional secretion of proteins. Two different definitions of context were evaluated: structural order and charge signature in the neighborhood of DE as one, and the nature of the amino acid insertion (‘X’) when a D-X-E motif is present, as another. We observe that among the eukaryotic proteins we studied, the odds of the protein being secreted are higher when the DE motif occurs in the disordered region of the protein, with higher hydrophobicity and lower charge or when the insertion X has a propensity for phosphorylation.

## 1. INTRODUCTION

Proteins need to be secreted outside the cell for several physiologically important reasons [1–3]. In eukaryotes, proteins with an N-terminal signal peptide are known to get conventionally secreted in a vesicular mode through endoplasmic reticulum (ER) – Golgi secretory pathway (ER-Golgi-secretory vesicles) [4, 5]. However, interestingly, many proteins which lack in such well defined signal peptides are also secreted, mostly under cellular stress [6]. This important class of unconventional protein secretion (UPS) reflects the cellular response to stressors such as inflammation, nutrient stress, ER stress, mechanical stress, etc and continues to grow in relevance as many such instances are detected in disease associated with dysfunctional autophagy such as neurodegeneration [7]. Secretion of leaderless proteins, without signal peptides, is intriguing and this non-canonical export mechanism raises several mechanistic questions on their presumably unique secretory pathways [6]. While at least four different UPS mechanisms have been identified so far [6, 8], even simpler and fundamental questions on how these leaderless proteins are identified remain open.

Conventional signal peptides are 15-50 amino acid tags (“zip code”) at the N-terminus of the proteins [4, 5, 9]. These signal peptides have a characteristic tripartite structure – positively charged N-region, a hydrophobic H-region and a polar C-region – which makes it easier for the export machinery in the cells as well as for the computational models, to sort the secretory from the nonsecretory proteins. Several predictive models for identifying these signals have been developed [10, 11]. On the other hand, identifying unconventionally secreted proteins has not been equally intuitive, as they lack in the pattern of canonical leader sequence [4, 12]. Modelling attempts to predict the unconventional secretion had limited success so far in deciphering the signatures of non-canonical secretion[13]. Using artificial neural networks which work recognizing with implicit patterns in protein sequences, unconventional secretory proteins were categorized by the properties such as amino acid composition, secondary structure and disordered regions but do not explicitly reveal the patterns among the unconventional secretory proteins [14, 15]. A rational basis for how non-classically secreted proteins are identified has been missing thus far.

In a recent landmark work, a motif which drives a protein through the non-canonical secretion pathway was identified for the first time. By systematically comparing the superoxide dismutase 1 (SOD1) from human and mouse cells with their extracellular SOD1 counterparts from human, mouse and yeast cells, a diacidic motif (asp-glu or D-E) had been identified to be responsible for its unconventional secretion. Of the 6 amino acid insertion in SOD1, compared to its extracellular homologs, UPS was found to be sensitive to the mutation of two amino acids aspartate (D) and glutamate (E) [16]. As a first and concrete observation, this finding raises the possibility that DE could be a generic UPS signal motif. However, as the short two amino acid length motif could occur in most proteins, in this study, we nucleate a minimal but rational hypothesis for the role of DE and its structural and charge context to act as a trigger for UPS.

## 2. MATERIALS AND METHODS

<u>Data curation</u> – All the proteins used in this analysis are eukaryotic proteins. Since the hypothesis was centered on the contexts in which DE acts as a UPS trigger, any protein which had no DE motif was discarded from the analysis. The proteins used in the analysis were chosen from three different groups:- Group 1: Since it was recently demonstrated [16] that the presence of DE in UPS of SOD1 (cargo involved in neurodegenerative disease, ALS), we intended to check the same in all other neurodegeneration causing aggregates such as α-synuclein, β-amyloid, TDP-43, Tau followed by Group 2 that encompasses all the known UPS cargoes known till date. All these secretory proteins are chosen from [17]. Group 3: Non–secretory proteins (cytoplasmic) were chosen from the human protein atlas (http://www.proteinatlas.org/humancell/cytosol). While the list of non-secretory proteins could be very large, we did not add any more of them than the number of secretory proteins to keep the data set groups unbiased. The complete list of proteins used in the analysis is tabulated in Table 1. The final curated set included 60 different eukaryotic proteins, 49 of which had both the DE motif as well as the structural data. 33 of these are non-secretory and 27 are secreted by unconventional protein secretion.

**Table 1:**
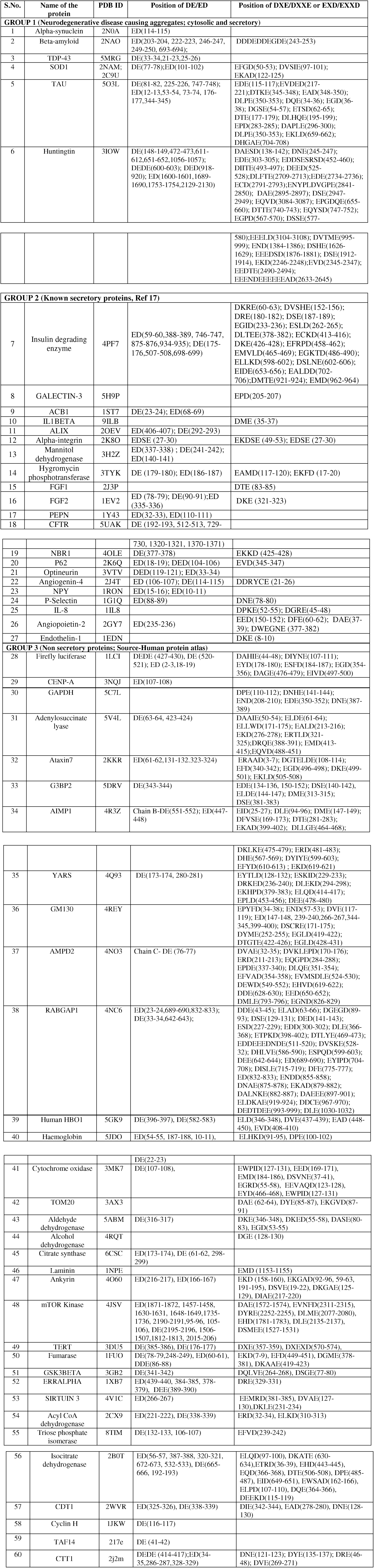
Eukaryotic proteins used in the present analysis: 60 proteins were curated and sorted as the ones secreted by unconventional secretion and those not secreted. The occurrences of DE/ED motif as well as insertions DXE/DXXE/EXD/EXXD are indicated.

<u>Variables describing the context</u>: For each of the selected proteins, three parameters defining the context in which DE occurs in a protein were defined: (1) Hydrophobicity (H) of the local region, by adding the hydrophobicity score of 3 amino acids on either side of the DE motif (2) Charge (C) of the local region, by summing the charge of 3 amino acids flanking on either side of DE and (3) structural order - the local structure around DE was initially categorically classified as ordered (O) if DE is in α- helix or β- sheet, disordered (D) if DE is in a loop or disordered region (as illustrated in Figure 1). If DE occurred in the border (B) of structured/unstructured regions, given the limited sample size of the data, we considered the option of regrouping B either as O or as D. Of the two choices, border region classified as the disordered region was a better choice *a posteriori*, comparing the quality of multivariate regression model for the prediction of secretion. So, in all the reported analyses, we only used the two structural order categories D (representing both D and B), and O.

**Figure 1:**
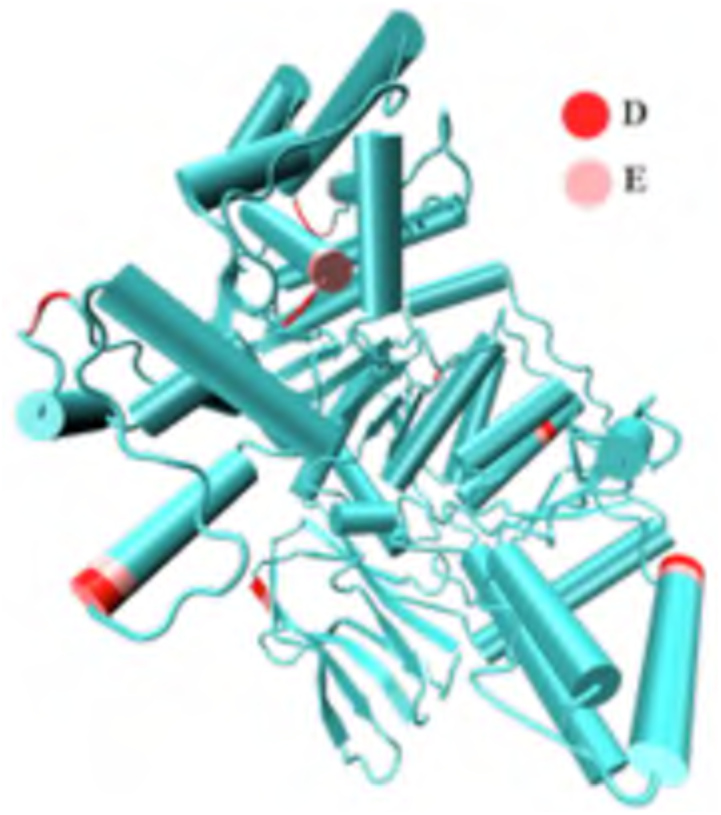
Illustration of the definition of structural order in NADP Isocitrate dehydrogenase (PDB: 2B0T). DE motif was seen in ordered, loop and border regions.

<u>Choosing the most relevant DE</u>: Many of the proteins we curated, secretory as well as nonsecretory, had multiple DE motifs. The working hypothesis in the present work is that of these multiple DEs, only one DE, depending on its context, is relevant for defining the UPS secretory fate of the protein and we identified this one DE with systematic screening. We created input files with all possible options, wherever there was an ambiguity: if the protein has DE in both disordered and ordered regions, one input data set which considers the protein as disordered and the other that considers it as ordered were prepared. The secretory predictions in the multivariate regression using either of these input files was used for deciding that if disorder and order are simultaneously present, it is considered to be disordered. Similar screening through different input files was used to reach the conclusion that when there are multiple DE motifs in disordered regions of the same protein, the DE that is present in the most hydrophobic or with least charge was relevant for UPS.

## 3. RESULTS

### 3.1 Nature of the neighbourhood of DE as a context

#### 3.1A Dependency on single variable

Before building a predictive model, the curated data was classified as secretory (UPS) and non-secretory and their relation to each of the individually chosen variables describing the structural context of where DE occurs in the protein was examined. The secretory nature has a dependence (Figure 2), although not very strong, on all three context variables hydrophobicity (H), charge (C), and (dis)order D/O that we calculated based on the protein sequence and structure data (**Materials and Methods** section).

**Figure 2:**
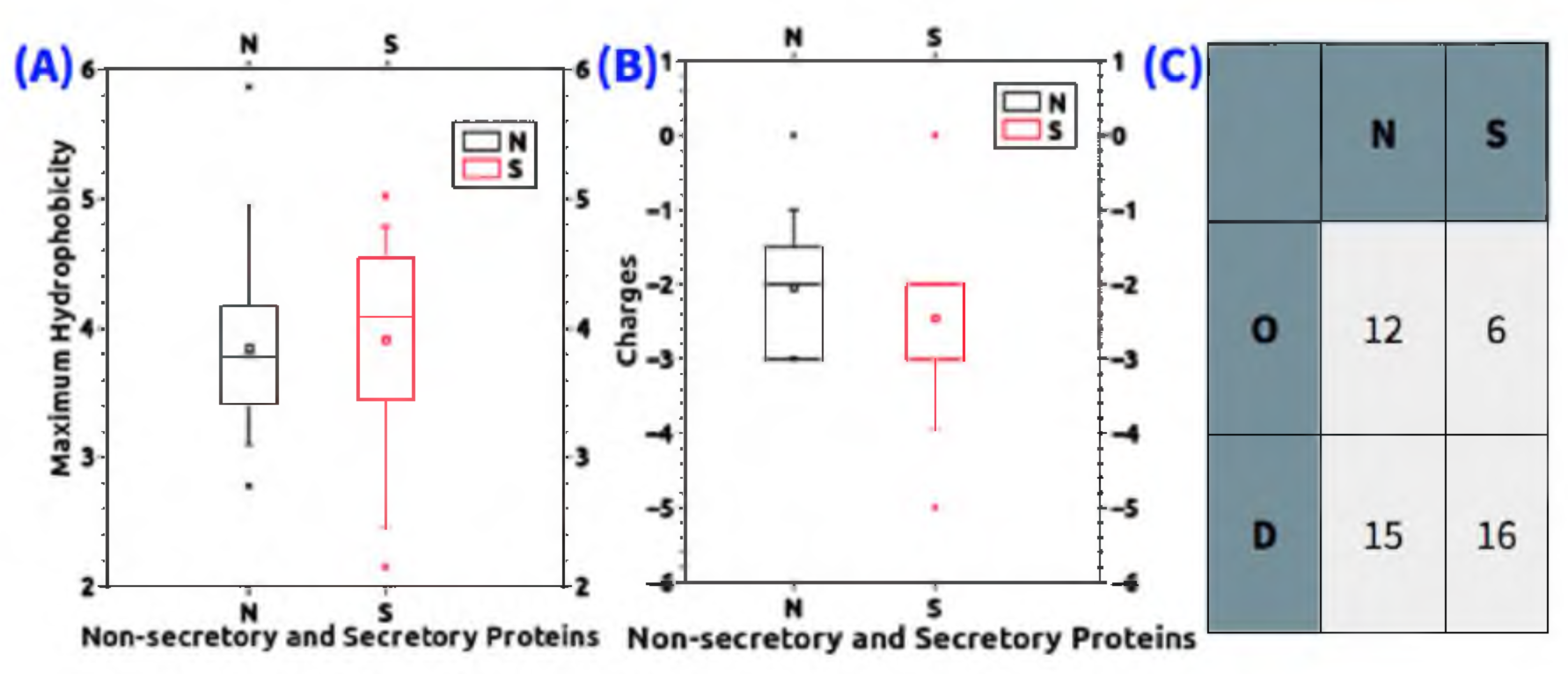
Depedence of secretory nature on the local properties of DE motif (A) hydrophobicity (B) charge (C) order.

#### 3.1B Multivariate regression

Since each of the three local parameters showed a relation to the secretory nature, we combined all of them in a binomial logistic regression model to predict the odds of secretion. The results summarized in Table 2 show that the odds of secretion are higher if the DE is in the disordered region, with higher hydrophobicity and lower charge. Among the multiple occurrences of DE in the protein, the results reported are the best predictions obtained by repeating the regression with the different DE’s and selecting the DE from the disordered, higher hydrophobicity (**Materials and Methods** section).

**Table 2:** Predictions of secretion from multi-variate logistic regression using 3 variables: The odds of secretion ( and significiance) of these variables are: Disorder 2.4 (p=0.17), Hydrophobicity 1.7 (p=0.24), Charge 0.5 (p=0.08).

|  |  | Prediction |  | Percent of correct prediction |
| --- | --- | --- | --- | --- |
|  |  | Secreted | Not-secreted |  |
| **Experimental observation** | Secreted | 11 | 11 | 50 % |
|  | Not-Secreted | 8 | 19 | 70 % |

#### 3.1C Molecular Dynamics Simulation

We also performed molecular dynamics simulation on the wild type SOD1 and a mutant with four amino acids simultaneously changed to alanines (G72A/P74A/R79A/V81A). The all-atom simulations were performed for 10 ns each. From the calculations we see no notable difference in the flexibility of the loop with the alanine mutations (Figure 3). Since the alanine scan mutations of the amino acids flanking DE did not affect the structural order or increase the charge, according to the proposed hypothesis these mutations need not reduce the chances of UPS. Indeed, this was the observation that mutations to alanine [16] did not affect UPS. While this does not prove the hypothesis, it serves as a consistency check and lays out the possibility of trying alternative structural order inducing mutations.

**Figure 3:**
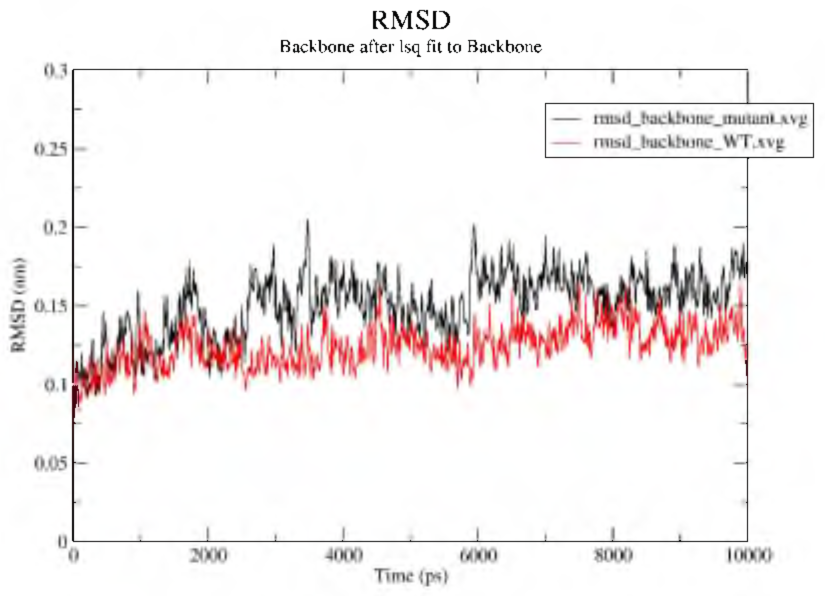
Results from classical all atom molecular dynamics o f SOD1 protein (PDB2C9U) performed using GROMACS with AMBER99SB forcefields. The root mean square deviation, which indicates the flexibility of the loop, was calculated relative to a reference structure from wild type and the mutant and was found to be comparable.

### 3.2 Amino acid insertion (D-X-E) as an alternative context

As a generalization of the context, we also examined the insertions X, among the occurrences of the D-X-E motif. D-X-E motif in the cytoplasmic domain of the membrane proteins has been implicated as a signal for the cargo transport [18]. It raises questions on whether the same motif could be relevant in other proteins too, especially because D-X-E has been implicated in the cargo concentration during the protein export from ER. There are several occurrences of D-X-E or D-X-X-E in our curated data, both in the secretory and non-secretory proteins. However, interestingly among the unconventionally secreted proteins, we observed the occurrence of DXE with one of the following recurring six amino acids that have the propensity for phosphorylation :- three O-phosphorylated amino acids (S, T, Y) or three N-phosphorylated amino acids (H,R,K) [19, 20]. The contingency table for these insertions and the possibility of secretion are mentioned in Table 3 which seems to be significant.

**Table 3:** Propensity of the amino acid insertion at DXE sites among secretory and non-secretory proteins: Among the secretory and nonsecretory proteins with a DXE motif, we analyzed to see how often X is one of the three O-phoshporylated amino acids (S, T, Y) or one of the three N-phosphorylated amino acids (H, R, K). The Fisher’s exact test for this contingency table gives a p-value < 0.01.

| **D-X-E insertion** | **Insertion**<br>**X=S,T,Y,H,R,K** | **Other insertions** |
| --- | --- | --- |
| Secreted | 7 | 1 |
| Unsecreted | 3 | 17 |

## 4. DISCUSSION

The role of DE in signalling had been noted in several other contexts [21]. For example, it had long been known that KDEL or HDEL sequences at the C-terminus act as retention signal to keep the proteins in ER [21]. Further, D-X-E was seen to be the signal for the export of transmembrane proteins such as VSV-G by COPII-coated vesicles from the endoplasmic reticulum to Golgi [18]. Although this pathway is mediated by the machinery (Sar1, Sec23/24) involved in the conventional secretory pathway, both these roles are different from the role of DE in SOD1 which is for unconventional protein secretion [16]. Thus, it is clear that the same DE motif depending on its flanking residues and insertions can assume a different signalling role. In this work, we examine several eukaryotic proteins to find clues for the patterns leading to UPS.

The aim of this work is not to predict the possibility of secretion by unconventional pathways but rather to build a hypothesis which may be accepted or rejected as and when more data will be available. Our preliminary analyses suggest that the diacidic motif DE when it appears in a disordered structural environment, with lower charge and higher hydrophobicity is likely to increase the odds of unconventional protein secretion. The p-values in the analysis were not significant at this stage, as the sample size of eukaryotic proteins with UPS is small. There are two ways to validate the hypothesis. The first is to curate data on more eukaryotic proteins which certainly go through UPS and check if the statistics improve significantly. Alternatively, since the hypothesis is centred around the structural order and charge on the amino acids flanking DE, SOD1 [16] secretion may be re-examined with mutations which promotes structural order or by increasing the charges, instead of substitutions to alanine as shown in [16].

Considering the alternative hypothesis proposed in this work, that D-X-E motif could be a signatory sequence for UPS, it is plausible that the post-translational modifictions, especially, phosphorylation of the inserted amino acids may act as a signal for UPS. It was observed that among the curated data the amino acids with propensity for O- or N-phosphorylation are inserted as DXE in the proteins undergo UPS. Contrary to this, the D-X-E motif that was seen in COPII is with aliphatic amino acid insertions [18]. Serine, threonine, tyrosine, as well as arginine [22], histidine and lysine [20] can undergo phosphorylation under oxidative stress, which is also known to be the external cue that triggers UPS. This hypothesis can be validated by searching for similar patterns in a larger dataset as well as by introducing the DXE motif and observing its chances for unconventional protein secretion.

## CONCLUSIONS

In this work, we introduce a hypothesis that the context of DE, defined by the structural order, charge and hydrophobicity around it, or an inserted amino acid which can easily undergo posttranslational modification may be a useful way of predicting whether the protein will undergo unconventional secretion. The size of the data on which these conclusions are drawn is small, as the scope was restricted to eukaryotic proteins known to be secreted *via* UPS. Despite this limitation, we find the conditions for the odds of unconventional secretion to be better. The hypothesis about structural order flanking DE or about the phosphorylation of the insertion DXE can be validated either with larger data sets or with experimental proof. With this study, we hope to raise testable hypothesis about the recognition of proteins secreted by unconventional pathways, something which remained underexplored as yet.

